# Muscle Diffraction at the Life Science X-ray Scattering Beamline

**DOI:** 10.64898/2026.02.11.705260

**Authors:** Khoi D. Nguyen, Anthony L. Hessel, Rachel L. Sadler, Nichlas M. Engels, Christine E. Delligatti, Samantha P. Harris, Lin Yang

## Abstract

We report on recent methodological advances at the Life Science X-ray Scattering (LiX) beamline of the National Synchrotron Light Source II (NSLS-II) to support small-angle X-ray scattering experiments on skeletal and cardiac muscle tissues. These experiments have been routinely performed at the BioCAT beamline of the Advanced Photon Source (APS) over the past two decades to measure sarcomeric protein organization within healthy and diseased muscle tissues and provide direct molecular evidence for their functional roles and dynamics. Many recent advances in our understanding of sarcomeric proteins relied on diffraction data and include, as examples, MyBP-C, crossbridge SRX/DRX states, and titin. With LiX now available for muscle experimentation, more muscle users can be supported which will speed up research of sarcomeric proteins, muscle biomechanics, and skeletal and cardiac myopathies. LiX explicitly focuses on high-throughput muscle diffraction with rapid sample turnover and semi-automated data processing. These operations have been tested and validated on skeletal and cardiac tissues sourced from both humans and multiple animal models including pig, rat, mouse, and zebrafish.

## Introduction

In the same way that laser diffraction applies visible light to measure inter-sarcomeric spacings of striated muscles at micrometer scales, X-ray diffraction applies light with angstrom wavelengths to measure intra-sarcomeric spacings of striated muscles at nanometer scales. These intra-sarcomeric spacings result from the semi-crystalline order of sarcomeric proteins, namely of actin, myosin, and regulatory proteins that attach to them and ultimately form the biomolecular basis for all muscle phenonmena (see Figure 1a-1b) (Ma and Irving, 2022; Reconditi, 2006; Brunello and Fusi, 2024). Historically, muscle X-ray diffraction methods were based on lab-based X-ray sources to identify the actin and myosin helical arrangements and hexagonal lattice order using long exposure (*∼*hours) of intact frog sartorius muscles, known for their excellent diffraction quality (Huxley and Brown, 1967). With the advent of synchrotron light sources providing X-rays with at least one million times more photon flux, these diffraction techniques push the envelope of ever shorter exposures (*∼*milliseconds) and higher signal strength (Huxley, 2004; Iwamoto, 2019; Squire, 2016). This allows for experimentation on a broader range of model organisms, time-resolved dynamics, smaller fibers, permeabilized preparations, and more difficult fibers sourced from myopathies that naturally diffract weakly.

**Figure 1.**
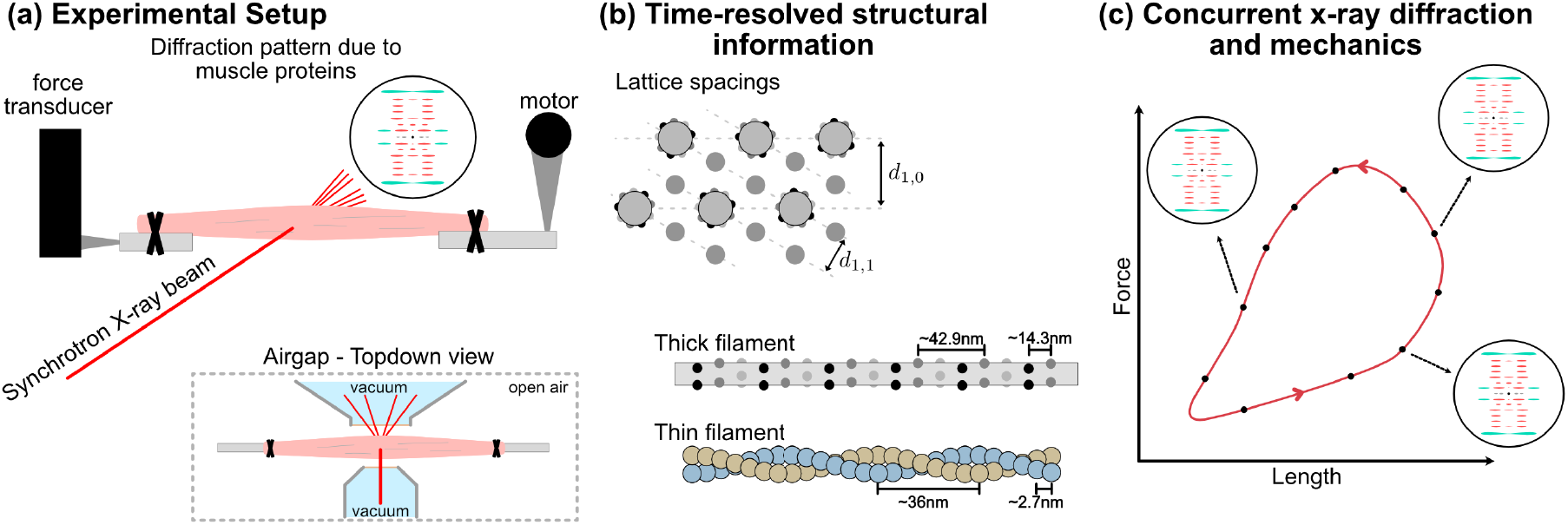
The muscle X-ray diffraction pipeline provides structural data regarding muscle protein organization at nanometer scales, often in conjunction with mechanics by coupling with force-length measurements. (a) The typical muscle mounting configuration with a motorized length lever and force transducer on a beamline airgap. The muscle can either be intact or permeabilized and range in size from single myofibrils to whole muscles depending on experimental design. (b) Common structural measurements include the two unit distances (d10 and d11) of the hexagonal grid formed by thick and thin filaments and the helical repeats of the filaments themselves. (c) Concurrent X-ray and force/length measurements provide nanomater-scaled structural information at different mechanical states.

Nowadays, synchrotron-based muscle X-ray diffraction is increasingly used as a core component of the muscle biologist’s experimental toolbox, and often in conjunction with mechanical experiments, to directly measure sarcomeric protein organization of both skeletal and cardiac muscle fibers under near-physiological conditions (Ma and Irving, 2022; Brunello and Fusi, 2024; Squire and Knupp, 2021; Powers et al., 2021; Iwamoto, 2019; Kuehn et al., 2025; Hill et al., 2025). Within the diffraction patterns are features termed equatorial and meridional reflections that are respectively normal to and in-line with the fiber’s long axis. Equatorial reflections result from structural protein organization along the cross-sectional plane of muscles fibers, and measures the hexagonal lattice spacings formed by interlacing thick and thin filaments and the relative distribution of electron density (or approximately mass) between components associated with the filaments (Malinchik and Yu, 1995). Equatorials thereby provide a quantifiable proxy for crossbridge attachment/detachment based on mass transfer between filaments. Meridional reflections on the other hand result from the structural protein organization in planes parallel to the fiber, and reveals information regarding filament strains and localization of different proteins to sarcomeric regions (Ma and Irving, 2022; Reconditi, 2006). Both equatorial and meridional reflections can be dynamically tracked during experiments and compared across perturbative conditions (e.g. pre/post drug incubation, healthy/diseased sample, and stretched/shortening cycles) to infer protein function, drug efficacy and toxicity, disease pathology, or otherwise evaluate muscle biomolecular states based solely on structural measurements.

Most forms of mechanics experiments on excised muscles can be adapted to fit into a synchrotron laboratory (termed beamline) configured for biological tissue processing (see Figure 1a). The technical specifications required for muscles include, from our experience, a q-range that spans molecular spacings as small as 5nm and as large as 50nm but broader ranges allow for more detailed muscle diffraction data, an adjustable beam spot size on the order of *∼*100µm, minimal photon flux of 1e10 photons/sec at the sample, and an open-air sample stage termed an airgap. The airgap is often designed to be as compact as possible with a width preferably on the order of centimeters since it is a tradeoff between the facts that X-ray beams must propagate in vacuum to avoid scattering by air molecules and that biological tissues cannot survive in vacuum. In terms of muscle hardware such as force transducers, motors, clamps, and electrodes, the challenge is to fit them all within the airgap while ensuring that the X-ray beampath is free and clear of all obstacles except for the muscle tissue itself.

Synchrotron-based muscle experiments in principle allow for concurrent measurements of muscle state based on sarcomeric protein organization along with mechanics (see Figure 1c). That structure-and-function relationships can be simultaneously determined and correlated is a powerful tool and adds a dimension of analysis to study and better model muscle phenomena under active research such as but not limited to residual force enhancement/depression (Hessel et al., 2024b; Joumaa et al., 2021), work loop dynamics (Pearson et al., 2025), activation/deactivation dynamics (Gong et al., 2022; Hill et al., 2022), Frank-Starling Law (Reconditi et al., 2017), spontaneous oscillatory contractions (Kono et al., 2020), and drug discovery applications (Pearson et al., 2025; Gollapudi et al., 2021). Yet synchrotron-based experiments are in general not accessible for the majority of muscle biologists owing to several major barriers of entry. First and foremost, one needs access to a beamline configured for muscles of which there are a handful worldwide. Until recently, there was only one heavily used muscle-capable beamline in the US, the BioCAT beamline at the Advanced Photon Source of Argonne National Laboratory. Access is secured through a months-long and competitive user proposal system typically in the form of a 24-hour or 72-hour continuous timeslot (termed beamline runs). Second, the prerequisite engineering and physics background needed to understand beamline operations, troubleshoot errors, and analyze the data form a steep but not insurmountable learning curve for most. Third, beamline runs are resource-intensive, high-pressured environments with little room for error where users typically work non-stop through the entire allocated beamtime to maximize productive value.

Here, we present methodological advances at the Life Science X-ray Scattering (LiX) beamline of the National Synchrotron Light Source II (NSLS-II) at Brookhaven National Laboratory based out of Long Island, NY to run high-throughput muscle X-ray diffraction experiments in a “plug-and-play” configuration. The express intent of these advances was to lower the barriers of entry for synchrotron-based diffraction experiments for muscle biologists who are not specialist synchrotron users themselves. LiX is designed primarily for biological sample testing and we refer to DiFabio et al. (2016) and Yang et al. (2022) for a detailed breakdown of LiX hardware and technical capabilities. Briefly, LiX is equipped with both small-angle and wide-angle X-ray scattering (SAXS/WAXS) detectors, an adjustable airgap to minimize background scattering due to air, variable beam spot size (5-500µm), and motorized sample stages. The NSLS-II facility itself is also equipped with state-of-the-art biosafety level 2 wet lab and animal care facilities available to users. We focused on upgrades that maximize throughput of common muscle diffraction experiments while limiting user-errors and without compromising on data quality. Through an iterative approach, we custom built a mobile modular experimental rig that can be configured with length controllers, force transducers, electrodes, solution exchangers, and temperature control as needed. Importantly, this rig is also designed with a rapid mount/dismount mechanism onto and off the beamline’s sample stage while keeping the muscle sample loaded. This not only allows for fast sample exchanges but also for running time-consuming experimental steps (e.g. sample loading and drug incubation) when dismounted. Thus, by running multiple rigs concurrently and mounting them onto the beamline only when needed, several muscle samples are processed in parallel which increases experimental throughput by several folds and makes optimal use of limited beamtime.

## Methods

The BioCAT beamline at the Advanced Photon Source (APS) has spearheaded the development of muscle X-ray diffraction technology over the past two decades through consistent support from the National Institutes of Health and has stood as the sole beamline in the US equipped for muscle experimentation (Irving et al., 2000). The methodological advances at LiX described here were developed in communication with BioCAT team and differentiated through an explicit focus on high through-put sample processing and a ground-up redesign of hardware, software, and data analysis. In this manner, muscle diffraction at LiX focuses on efficiently running fiber experiments already validated at BioCAT and do not necessarily require its bleeding-edge muscle diffraction capabilities.

### LiX configuration

The beamline accommodates high-throughput experiments on biological samples and runs a variety of experiments across multiple fields (DiFabio et al., 2016; Yang et al., 2020, 2021, 2022; Yang, 2024). The spot size (5 - 500µm) and X-ray energy (7-18 keV) can be adjusted based on experimental needs. Furthermore, it is equipped with simultaneous data acquisition on wide-angle (Pilatus3 900k) and small-angle (Pilatus3 1M) detectors, whose distance from the sample can be set to cover the desired q-range. For muscle diffraction specifically, the beamline operates under a scanning configuration at *∼*15keV, a photon flux of *∼* 5e10 photons/sec at the sample, and detectors positioned to cover the entire q-range from 0.005^°^A^-1^ to 2^°^A^-1^. The beam spot size is optimized for the muscle type and experiments at hand and typically lies between 25 and 100µm at the sample by controlling the beam’s divergence through a secondary source aperture. Samples are mounted on an open-air sample stage termed an airgap that is adjusted to be as compact as the mounting hardware allows (*∼*2cm) to minimize signal degradation due to air scattering. A camera placed in-line with the X-ray beam provides precise spatial positioning of the muscle sample relative to the beam.

### Muscle X-ray diffraction hardware

Although the X-ray component is crucial, muscle samples require only a few minutes on the beamline to image a diffraction pattern, a minor fraction of time when compared to other steps such as sample loading, calibration, and incubation times. Thus to maximize throughput without compromising on rigor or data quality, we optimized the hardware at LiX to run multiple muscle mechanics experiments in parallel. We built a stand-alone muscle mechanics rig that can be mounted onto the beamline with a preloaded muscle sample only when needed for X-ray exposure, and then quickly dismounted and relocated to a lab bench for all other steps. In this manner, deadtime at the beamline is minimized and throughput is limited only by skilled hands available to run the experiments. At our current operational capacity, we run 5 experiments concurrently with roughly 3 minutes of deadtime between imaging diffraction patterns. We designed the muscle mechanics rig as a mobile, stand-alone modular unit with a rapid mount/dismount mechanism and with equipment/computers attached to a portable power station to ensure continuous power during transport to and from the beamline. These rigs are shown in Figure 2. The modularity is to accommodate the wide range of muscle experiments by allowing for piecewise selection of equipment as needed such as force transducers, muscle length motors, electrodes, temperature control, solution exchangers, oxygen bubblers, and equipment otherwise not considered here. The mobility and mounting mechanism are based on optomechanical parts and allow for rapid sample interchanges and sample manipulation off the beamline. The muscle samples themselves are attached to two separate 3-axis linear micromanipulators via specialized clamps made from hypodermic needles. Other methods of attachments such as T-clips, hooks, sutures, and glue will also work with minor modifications. When mounted to the beamline, muscle samples rest inside a temperature-controlled solution bath coupled to a motorized z-stage that moves it out of beampath during data acquisition but leaves the muscle unperturbed. This ensures that the diffraction pattern is imaged while the muscle is held in-air and avoids signal degradation largely due to absorption by the buffer solution. Care is also taken to ensure that the muscle sample is in-air for no longer than 20s. Multiple temperature controlled solution baths are available and can be quickly exchanged to switch between solution types as needed based on experimental design. For muscle samples that require continuous exposure to solution (e.g. actively contracting intact muscles), the diffraction pattern is acquired with the solution bath still in place. To accommodate these situations, we designed the solution baths with a narrow form factor and thin Kapton windows so that the X-ray beam travels through as little buffer solution as possible.

**Figure 2.**
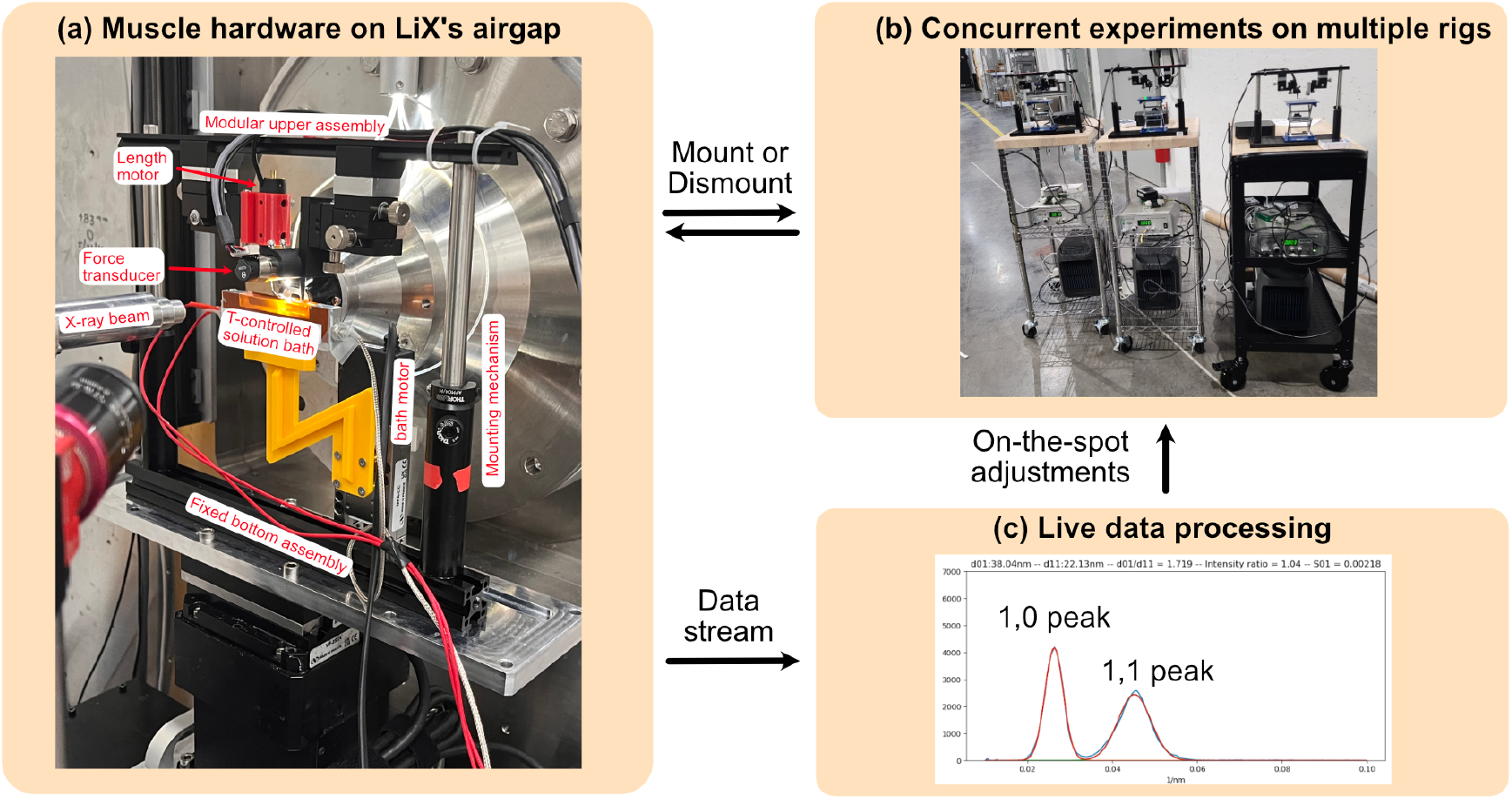
The high-throughput MyoSAXS setup at the Life Science X-ray Scattering (LiX) beamline of the National Synchrotron Light Source II. (a) The experimental rig consisting of a fixed bottom assembly, a dismountable upper assembly containing the muscle sample, and a temperature-controlled solution bath that moves out of the beampath during data acquisition to avoid solution scattering. (b) The upper assembly consists of length controllers and force transducers for muscle mechanics and mounts onto fully self-contained mobile rigs for time-intensive solution incubations and sample changes outside the hutch. This allows for running multiple rigs simultaneously which increases experimental throughput. (c) Diffraction patterns are streamed and analyzed in real-time to allow for on-the-spot quality checks and adjustment of experimental protocol as needed.

### Data acquisition and processing

Imaging a muscle sample’s diffraction pattern follows one of several approaches. The first is to identify a single high-quality spot by running an initial pass at very low X-ray exposures across a wide area, and then imaging the selected spot multiple times across different experiment conditions at higher X-ray exposures. This provides exceptionally clean diffraction patterns but risks tissue damage due to repeated X-ray exposures and is best used for experimental designs with single or very limited number of conditions. Another limitation with spot selection is that the spot is often in a similar part across all samples tested, and the data therefore does not provide a spatially average representation. A second approach is to scan a large portion, if the not entirety, of the muscle while under continuous X-ray exposure. This operation generates a single or few diffraction patterns and is effectively a spatial low-pass filter on the entire muscle sample. Exceedingly long X-ray exposures without tissue damage are possible here because no single region is overexposed and the result is diffraction patterns with excellent signal-to-noise ratio. The major drawback is an imaging artifact similar to motion-blur because spatially heterogeneous parts of muscle samples are all superimposed on each other, and thus this approach is best used for homogeneous muscle samples known to have long-range spatial order or for weakly diffracting muscle samples that necessitate long exposures.

To optimize the benefits of both spot selection and scanning imaging approaches, we take a third approach at LiX that acquires many pseudo-spot images with short exposures as the muscle is in motion in a raster area scan. Using a beam spot size (25 - 100µm) that is small in comparison to muscle samples, typically several millimeters in length and several hundred microns in diameter, and exposures ranging from 100ms to 1s, each muscle sample generates dozens-to-hundreds of images that are merged together in post-processing to form one final merged image per muscle sample per condition. The post-processing pipeline is fully automated using custom python scripts that correct each image for fiber orientation (fibers within a bundle are not always in perfect alignment), filters out images with low signal-to-noise ratio, and remove images in regions with mostly connective tissue. This approach adds granularity to the data acquisition to accommodate spatial heterogeneity of muscle samples due to connective tissues, misaligned or crossed fibers, or regions damaged during preparation that diffract poorly, all of which are removed prior to merging. Although data-intensive and computationally heavy, this approach of raster scanning a large area avoids both tissue damage due to overexposure and motion-blur-like imaging artifacts.

The merged images are then semi-autonomously analyzed to provide measurements of lattice space based on equatorial 1,0 and 1,1 reflections, myosin head placement based on equatorial intensity ratio *I*_1,1_*/I*_1,0_, and myosin and actin repeats along the filaments based on their nth order meridional reflections. This step emulates much of what MuscleX, an open-sourced suite of data tools commonly used by users to process muscle diffraction data (Jiratrakanvong et al., 2024), does, but optimized for LiX data structures and real-time analysis. It requires an initial calibration process in terms of selecting background subtraction parameters and reasonable bounds for Gaussian curve-fitting. We note here that these parameters vary depending on muscle types, experimental conditions, and tissue quality and so some curation and therefore user-bias is currently needed. The entire processing pipeline is performed initially in real-time using custom python scripts on LiX servers to enable on-the-spot adjustments as needed such as scanning a different area on the sample, tuning exposure settings, or modifying the experimental protocol to maximize diffraction quality. Real-time data analysis and the ability to make on-the-spot adjustments maximize data quality and avoid repeat experiments due to flaws normally only found in post-experiment data analysis.

### Muscle study of fMyBP-C

To demonstrate the technical capabilities and throughput of LiX, we ran an archetype experimental design in which diffraction patterns taken before and after a treatment are compared. In this case, the treatment is of a previously defined effect from a BioCAT study focused on the role of fast myosin-binding protein C (fMyBP-C) in mediating myosin dynamic forces of skeletal muscles via the genetically modified mouse line SnoopC2, which was specifically engineered to controllably cleave away N-terminal fMyBP-C by incubation with tobacco etch virus protease (TEVp) (Hessel et al., 2024a). Using permeabilized fiber bundle preparations of psoas muscles, Hessel et al. (2024a) found that fMyBP-C cleavage expanded the actomyosin lattice, stretched the filaments themselves, and shifted more myosin heads from the thick filament to thin filaments. Here, we reproduced the experiment at LiX using extensor digitorum longus (EDL) permeabilized fibers prepared from SnoopC2 mice. Both EDL and psoas muscles are *>*90% fast twitch in mouse (Song et al., 2021; Baby et al., 2025) and thus we expect EDL to show a similar response to psoas upon fMyBP-C cleavage in terms of directionality but potentially different in absolute value or magnitude. We went one step further beyond the referenced study by introducing myosin deactivation agent mavacamten after fMyBP-C cleavage to investigate its therapeutic potential in reversing the known downstream effects of fMyBP-C removal. The effect of mavacamten on skeletal muscle myosin heads was also previously defined in another study at BioCAT (Kuehn et al., 2025).

The Hessel et al. (2024a) study at BioCAT on SnoopC2 psoas tissue consisted of *N* = 33 fiber bundles, held at 2.7µm sarcomere length (SL), room temperature, with X-ray diffraction data taken at two states in series (control, +TEVp). The study here at LiX on EDL tissue consists of *N* = 29 fiber bundles, also held at 2.7µm SL and room temperature, but instead consists of X-ray diffraction data taken at three states in series (control, +TEVp, and +mavacamten). The control state consists of fiber bundles imaged in relaxing solution without any added compounds. The +TEVp state is after incubation (100 units of acTEV protease per 600 µL relaxing solution) and imaged again in relaxing solution. The +mavacamten state is after a second incubation (50µM mavacamten in relaxing solution) and imaged a third time in the incubating solution. The BioCAT study was conducted over two separate 72-hour beamline runs spaced several months apart. The LiX study in comparison was completed in a single 72 -hour run, which speaks to the explicit emphasis on throughput for muscle diffraction experiments at the LiX beamline.

The statistical analysis consists of a mixed model ANOVA with main effect of condition (control / +TEVp / +mavacamten), and individual random effect to account for repeated measures across conditions. Response data was best Box-Cox transformed to meet normality assumptions. A significant main effect was followed up with Tukey’s highly Significant Different (HSD) post-hoc procedure for multiple comparisons with alpha set at 0.05.

## Results and Discussion

General quality of LiX datasets: With muscle diffraction now available, LiX has imaged several muscle types from animals ranging from zebrafish (Mead et al., 2024) to humans (Ochala et al., 2025) with diffraction quality strongly dependent on muscle type and disease state. Figure 3 shows a representative selection of permeabilized preparations from mouse and rat skeletal muscles and pig cardiac muscles under passive conditions. Certain reflections are readily resolvable in most muscle types and disease states. These include the principal equatorial reflections (1,0 and 1,1) that measure lattice spacings and relative mass between thick and thin filaments and meridional reflections (M3 and actin layer line 6) that measure the nth order reflections of the principal myosin and actin repeats. The diffraction quality of other reflections such as myosin layer lines, low-order M clusters, M6, M11, actin layer lines 5 and 7, higher order equatorials are all case-specific, each of which map to different features of thick and thin filaments (Koubassova et al., 2025; Ma and Irving, 2022; Brunello and Fusi, 2024; Reconditi, 2006). We have had successes in imaging these more difficult reflections at LiX in some muscle types with further optimizations in sample preparation protocol, beam spot size, and exposure settings but success across the board is not guaranteed. Thus, for those interested in these more-difficult reflections and for new users in general, we recommend initial pilot studies of a few samples at LiX to first optimize diffraction quality prior to committing to a full 72-hour beamrun.

**Figure 3.**
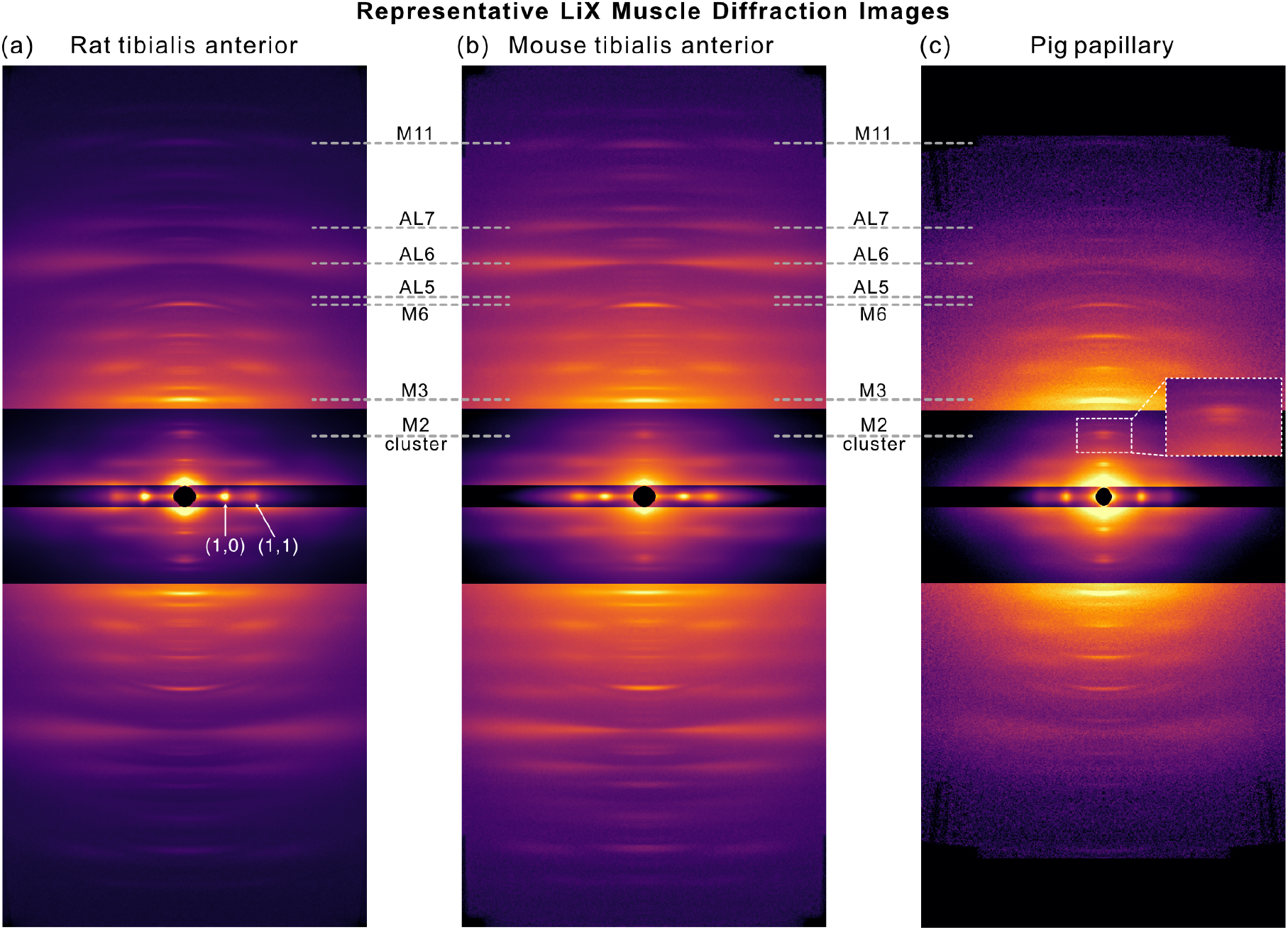
Representative diffraction patterns of muscle types imaged at LiX that include permeabilized preparations under passive conditions of (a) rat tibialis anterior, (b) mouse tibialis anterior, and (c) pig papillary tissue with inset showing M2-cluster resolving into individual peaks. Images are divided into different zones and arbitrarily scaled for better visualization. Common reflections of interest are annotated in the form of M# or AL# where M stands for myosin, AL for actin layer line, and number the nth order.

### Case study with the SnoopC2 mouse model

To stress test LiX’s muscle diffraction capabilities and throughput in a real use-case, we acquired a full dataset in a single 72-hour beamline run that investigated the downstream effects of fMyBP-C removal in the SnoopC2 mouse line engineered to cleave fMyBP-C upon addition of TEVp. The experimental design is similar to a prior dataset acquired at BioCAT that required two separate 72-hour beamline runs and thus provides a point of comparison in terms of data quality, consistency, and throughput. Figure 4 summarizes the dataset by visualizing shifts in muscle diffraction values, and in all cases shows that the LiX dataset recapitulates the same directionality and conclusions of fMyBP-C cleavage by TEVp as in the BioCAT dataset. Namely, fMyBP-C cleavage expands the actomyosin lattice (increased lattice spacing d10), allows more myosin heads to move to thin filaments (increased *I*_1,1_*/I*_1,0_ ratio), and stretches the thick filaments (increased M3 spacing). Furthermore, a second treatment using myosin deactivator mavacamten in the LiX dataset shows partial reversal of fMyBP-C cleavage effects where *I*_1,1_*/I*_1,0_ returns to pre-cleavage values, and d10 and M3 spacings only partially returns with values not different from either control or +TEVp states based on ANOVA testing. Taken together, this data suggests that mavacamten shifts myosin heads back to the thick filaments but not necessarily in the same molecular configuration as prior to fMyBP-C cleavage. Although the LiX and BioCAT datasets show the same directionality between control and +TEVp states, differences in magnitude between the two datasets are also significant and we attribute this change to different muscle types available to us at the time, i.e. psoas at BioCAT and EDL at LiX. Diffraction measurements are in principle agnostic to the originating beamline so we do not expect beam-specific settings such as photon flux, exposure times, and beam spot size to be significant contributors to the observed magnitude difference so long as signal-to-noise ratio is adequate. But more thorough investigations which systematically look into differences introduced by different beamlines are more than warranted, especially in cases where users are combining muscle datasets acquired from both LiX and BioCAT.

**Figure 4.**
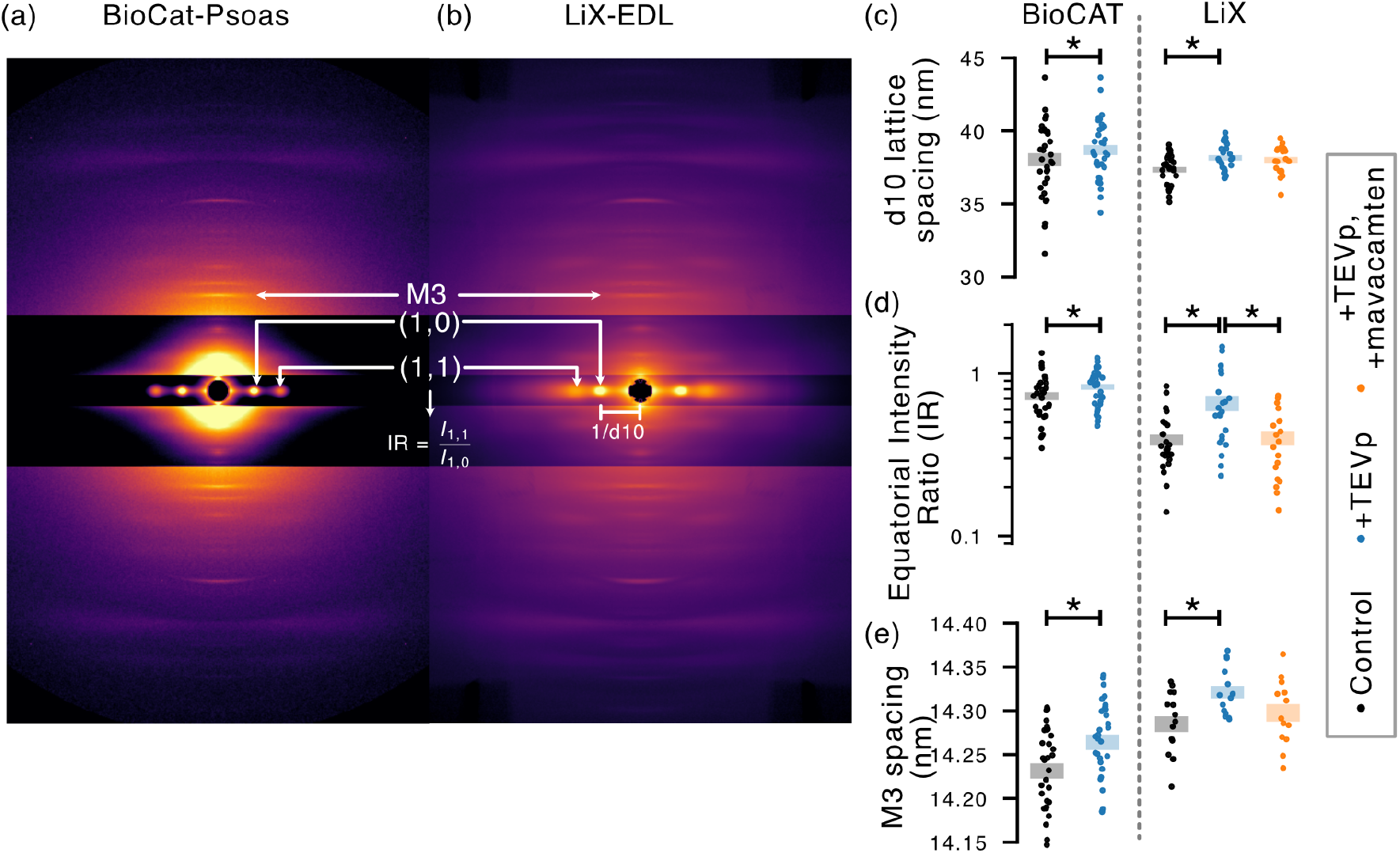
A muscle diffraction study on the downstream effects of fMyBP-C removal in mouse skeletal EDL muscles to stress test LiX’s muscle capabilities. The results of a similar study at BioCAT on mouse psoas muscles are shown for comparison. (a-b) Side-by-side comparison of representative image quality between BioCAT and LiX. Both images are control muscle samples at 2.7µm SL and room temperature. (c-e) Measurements of key diffraction features for control, fMyBP-C removal via TEVp cleavage, and partial reversal of fMyBP-C removal via the addition of myosin deactivator mavacemten in the LiX-EDL study. Lattice spacing d10 reflect the spacing between actomyosin filaments, intensity ratio *I*_1,1_*/I*_1,0_ the relative distribution of myosin heads between thick and thin filaments where higher values implies more crossbridge attachment, and M3 spacing the thick filament strains. Color bars indicate mean ± se and asterisk indicate statistically significant differences between groups (*p* < 0.05).

### Ongoing developments at LiX

We continue to improve the muscle diffraction capabilities at LiX on several fronts to further push on experimental throughput and complexity. The first is time-resolved experiments to study dynamic muscle phenomena such as tracking changes to sarcomere structures during dynamic length changes. The biggest limiting factor here is the X-ray exposure time required for sufficient signal-to-noise ratio. Our best efforts thus far consist of 33ms temporal resolution in intact preparations of rat left ventricle cardiac slices (unpublished data). BioCAT’s beam optimizations configuration for comparison has achieved sub-millisecond temporal resolution in certain use-cases. Thus, with the current muscle diffraction capabilities at LiX, time-resolved experiments with the highest likelihood of success are those involving muscle samples known to diffract strongly (e.g. intact preparations) and where slower dynamics (*∼*50ms or greater) are of interest (e.g. residual force enhancement/depression, slow oscillatory rheology, work loop analysis at *∼*3Hz or slower). The second development is inclusion of WAXS data analysis to extend the meridional axis beyond the normal SAXS q-range to investigate higher order myosin repeats (15^th^ order and beyond) and the principal actin repeat at *∼*2.7nm. Muscle diffraction studies have historically focused on low order reflections but recent studies have brought renewed interest to higher-order reflections to measure filament strains with higher sensitivity than possible in the SAXS q-range (Ma et al., 2018; Squire and Knupp, 2021) or to reconstruct filament structures at finer scales (Koubassova et al., 2025; Dutta et al., 2023). LiX is already configured for simultaneous SAXS/WAXS operations using multiple synchronized detectors but the WAXS data analysis for muscle diffraction remains to be developed and tested in a real use-case.

## Conclusions

Although X-ray diffraction experiments on living muscles have been around since the 1950s, it has historically remained a technically difficult experiment available to few due to high barriers of entry. These barriers include access to synchrotron radiation facilities, of which there are only a few in the US and around *∼*20 worldwide, and required technical expertise in engineering, optics, and beamline operations that presents a steep learning curve for most. The methodological advances presented here to support muscle diffraction experiments at LiX are geared towards lowering those barriers of entry. LiX now runs muscle diffraction experiments in addition to BioCAT and explicitly focuses on high-throughput experiments that do not necessarily require the bleeding-edge muscle diffraction capabilities of BioCAT, thereby expanding the total research capacity available for these experiments. The standardized hardware is designed to accommodate most forms of mechanics experiments (e.g. force-length, force-velocity, ramps, sinusoidals, and step perturbations) during simultaneous X-ray exposure and now provides a “plug-and-play” option for new users to run studies with minimal engineering effort. And lastly, the data acquisition and processing pipeline outputs result in real-time to help users troubleshoot on-the-spot as needed rather than on a return trip to LiX. These efforts, as a whole, develop the muscle X-ray diffraction pipeline into a more readily available technology for those whose research would benefit from the structural data it provides but previously could not access it.

## Acknowledgements

We thank Bradley Palmer and Matthew Caporizzo of University of Vermont for extended discussions, Nikoloas Papachatzis of Yale University for assistance with beamline operations, and the LiX beamline staff for their technical support.

## Data and Resource Availability

All relevant data and details of resources can be found within the article and its supplementary information.

## Author Contributions

KN, ALH, LY, SPH conceptualized the experiment. KN, ALH, LY designed the hardware, software, methodology and experiments. SPH provided the research material. KN and ALH supervised the experiments. KN wrote the original draft of the manuscript. All authors contributed to data collection, analysis, interpretation, and edited the final draft of the manuscript.

